# Monitoring soil microorganisms with community-level physiological profiles using Biolog EcoPlates™ in Chaohu lakeside wetland, east China

**DOI:** 10.1101/616821

**Authors:** Zhen Teng, Wei Fan, Huiling Wang, Xiaoqing Cao, Xiaoniu Xu

## Abstract

Under the circumstance of wetland degradation, we used Biolog EcoPlates™ method to investigate the impact of ecological restoration on the function of topsoil microbial communities by monitoring their metabolic diversity around Chaohu lakeside wetland. Four restoration patterns including reed shoaly land (RL), poplar plantation land (PL), abandoned shoaly grassland (GL) and cultivated flower land (FL) were selected. The result showed a rapid growth trend at the initial stage of incubation, following the fastest change rate at 72 h in both dormant and growing seasons, and the AWCD values of RL pattern was the highest at the detection points of each culture time, while the GL were the lowest. The calculation of diversity indicators also displayed significant lower McIntosh index in dormant season and Shannon-Wiener index in growing season in GL than in the others (*P* < 0.05). Carbohydrates and carboxylic acids were found to be the dominant substrates used in dormant season, whereas amino acids, polymers and phenolic acids were increasingly utilized by the microbial communities in growing season. We observed soil total potassium as the key factor that significantly affected the utilization efficiency of different carbon sources in both seasons (*P* < 0.05).

## Introduction

As one part of the terrestrial carbon pool, wetlands play a crucial role in global carbon cycling process [1]. Soil is the main component of the wetland ecosystem, and it can be strongly impacted by the hydrological changes caused by alternation of wetting and drying, which alter the edaphic redox environment and thus control biogeochemical processes [2]. With the development of urbanization, wetland environment has been degraded resulted from agricultural reclamation, global climate change and acid precipitation [3–5], which makes wetland become one of the most threatened ecosystems in the world [6]. Nowadays people’s awareness of ecological and environmental protection has been raised, and different restoration methods have been implemented to improve the wetland soil quality and species diversity. However, wetland carbon sequestration is still threatened by global warming [7]. It was reported that exogenous heavy metal inputs from agricultural development are introduced into natural wetlands [8, 9], which may significantly affect carbon cycling and balance of the area.

In a whole wetland ecosystem, microorganisms, as the decomposer and basic agent of the element circulation, play a key role in soil formation process [10, 11]. To make scienti fic researches in the reconstruction and restoration of urban lakeside wetland, it is necessary to explore soil microbial communities that affecting nutrient cycling and functional metabolism, especially for the utilization of carbon substrates. Although microorganisms are the important component for functioning of wetland ecosystem, their metabolic versatilities are poorly understood due to the complicated environmental stresses and regional microbial differentiation [12]. The Biolog EcoPlates™ is a method that relatively easy to operate, which is generally used to describe the diversity of community-level physiological profiles (CLPPs) [13–15]. Based on the biological and biochemical properties, the Biolog EcoPlates™ method can quickly characterize the ecological status of environmental samples [16], such as activated sludge [17], wastewater [18], sediments [19], and soils [13, 20].

As the fifth largest freshwater lake in China, Chaohu Lake wetland has been suffered from long term interference and damage due to the irrational use and excessive exploitation of resources in agricultural and construction activities [21–23]. Luckily, the environmental problems resulting from artificial disturbance has raised widely concern to the public. Native vegetation was basically non-existent, whereas since the year of 2003, large portions of the reclaimed land have been gradually restored to wetland and artificial forest [24, 25]. This is an extensively employed ecological restoration technique that plays a vital role in the Chaohu Lake watershed management [26]. Recent efforts have been done on the eutrophication of water ecosystem about Chaohu Lake [23, 27], however, there were little researches analyzing the functional metabolism shifts of microbial communities during the ecological restoration in this wetland ecosystem.

The objective of the present study was to investigate whether and how ecological restoration affect the wetland microbial community ecological functions along the Chaohu lakeside wetland. Based on the Biolog method, we further intended to find the specific metabolic characteristics and seasonal shifts of topsoil microbial communities in different patterns, which may help to estimate the efficiency of various restoration ways from the perspective of functional diversity of soil microorganisms.

## Materials and methods

### Site Description

This study was conducted around the northwest Chaohu Lake wetland between Pai River and Nanfei River, approximately 15 km from southern Hefei City, Anhui Province, China (30°25′28″N~31°43′28″N, 117°16′54″E~117°51′46″E). The area is influenced by the subtropical humid monsoon climate, and there are more than 230 days frost-free period, with the annual mean temperature and precipitation of 15-16°C and 900-1100 mm, respectively. The coastal soil types of Chaohu Lake and its main rivers belong to paddy soil.

### Sampling Design and Field Measurement

Four types of wetland soils were selected representing different restoration patterns, including reed shoaly land (RL, natural restoration pattern), poplar plantation land (PL, artificial restoration pattern), abandoned shoaly grassland (GL, Control) and cultivated flower land (FL, artificial restoration pattern). PL site was located at the Hefei Lakeside Wetland Forest Park, FL and GL sites were chosen from the Dawei Eco-agricultural Park, and RL site was selected randomly along the basin area, with specific marks on the sampling sites. Field measurement was implemented in the vegetation dormant and growing season (early March and late July 2018). Basic information and vegetation characteristics of the experimental sites were shown in Table 1.

**Table 1.** Basic characteristics of experimental sites in Chaohu lakeside wetland.

| Pattern | Latitude | Longitude | Dominant vegetation |  |  | Hydrological condition | Underground water level (cm) |
| --- | --- | --- | --- | --- | --- | --- | --- |
|  |  |  | Species | Family | Genus |  |  |
| GL | 31°40'43" | 117°17'37" | *Digitaria sanguinalis* | Poaceae | *Digitaria* | Seasonal flooding | 80 |
| RL | 31°43'14" | 117°21'58" | *Phragmites australis* | Poaceae | *Phragmites* | Seasonal flooding | 70 |
| PL | 31°43'27" | 117°23'22" | *Populus deltoides* cv. 'I-69';<br>*Populus euramevica* na cv. 'I-214' | Salicaceae | *Populus* | Seasonal flooding | 100 |
| FL | 31°39'55" | 117°15'52" | *Zinnia elegans* Jacq. | Compositae | *Zinnia* | Intermittent flooding | 80 |

After removing the litter layer, we collected several soil subsamples using a soil auger (6 cm in diameter), mixed together as one sample in each plot, and three plots were selected for biological replicates of each pattern. The samples were stored into sealed polyethylene bags and transported to the laboratory in a cooler box with ice bags. After passed through a 2 mm sieve to remove roots and other debris, the samples were divided into two equal parts. One part was for Biolog EcoPlates™ analysis, another for soil physiochemical property measurement, including soil water content (SWC), pH, dissolved organic carbon (DOC), ammonium nitrogen (NH_4_^+^-N), nitrate nitrogen (NO_3_^−^-N), soil organic carbon (SOC), total nitrogen (TN), total phosphorus (TP) and total potassium (TK) contents.

### Soil characteristics analysis

SWC was measured by oven-drying at 105 °C to a constant weight. Soil pH was determined using a pH meter in 1:2.5 soil / water suspensions. NH_4_^+^-N and NO_3_^−^-N were measured by a flow injection auto-analyzer (FIAStar 5000, FOSS, Sweden), as well as TP after micro-Kjeldahl digestion. SOC and TN were measured with a CN Analyzer (EA 3000, Vector, Italy). DOC was determined using a TOC auto-analyzer (Multi N/C 3100, Jena Analytik, Germany) [28]. In detail, one part samples (30 g fresh soil) were extracted with 50 mL of 0.5 mol·L^−1^ K_2_SO_4_ solution, shaking for 30 minutes and then filtered. The filtrate was diluted and determined by the TOC instrument. TK was measured by the Atomic Absorption Spectrophotometer (TAS-990AFG, Pgeneral, China) after micro-Kjeldahl digestion.

### Community Level Physiological Profiling

The Biolog EcoPlates™ method was used to conduct a 7-day dynamic monitoring on the functional diversity of soil microbial community, and the average well color development (AWCD) values of carbon-source utilization were collected [29]. Every EcoPlate had 96 wells containing 31 different carbon sources plus a blank well, in three replications. The utilization rate was pointed by the reduction of tetrazolium violet redox dye, which changed from colorless to purple if the single source was used by added microorganisms [16]. In addition, EcoPlate substrates were subdivided into six groups: amino acids, amines, carbohydrates, carboxylic acids, phenolic acids and polymers [30].

The EcoPlates to be tested were prepared in the following way: 10 g of fresh soil was weighed and put into a 250 mL triangle bottle. Then we added 100 mL sterilized 0.85% NaCl solution, shaking for 30 min (speed at 170 r·min^−1^). The turbid liquid was diluted with ultrapure water for 1000 times and incubated at 4°C for 2-3 min. The supernatant was taken into sterile culture dish and drew 150 µL per channel with the eight channel pipetting gun into the EcoPlates. Finally the plates were cultured on 28°C in biochemical incubator for 7 d. Absorbance at 590 and 750 nm was measured on Biolog Microstation after 24, 48, 72, 96, 120, 144 and 168 of incubation hours. Optical density (OD) value from each well was corrected by subtracting the control (blank well) values from each plate well. The OD values obtained at 72 h represented the optima range of optical density readings, so 72 h of incubation results was used for assessing the microbial functional diversity and statistical analyses.

### Statistical Analysis

In the process of Biolog data analysis, the AWCD value was calculated to reflect the overall activity of soil microorganisms [31], and the Shannon-wiener index (*H*′), McIntosh index (*U*) and Simpson index (*D*) at 72 h in the process of cultivation were investigated to represent the metabolic functional diversity of soil microbial community (Table 2)[32, 33].

**Table 2.** Formulae for calculations.

| Index | Definition | Formulae | Notes |
| --- | --- | --- | --- |
| Average well color development | / | $AWCD = \sum (C_i - R) / 31$ | $C_i$ is the OD value of each substrate containing well (590-750 nm). |
| Simpson index | Measure of evenness | $D = 1 - \sum P_i^2$ | $R$ is the OD value of the black well. |
| Shannon-Wiener index | Measure of richness | $H' = -\sum P_i \ln P_i$ | $P_i = (C_i - R) / \sum (C_i - R)$ |
| McIntosh index | Measure of diversity | $U = \sqrt{\sum n_i^2}$ | $n_i = (C_i - R)$ |

We performed an analysis of variance (ANOVA) using IBM SPSS 22.0 to explore the significant effect of the amendments on AWCD [16], functional diversity indices [33] and soil parameters, with the level of significance at *P* < 0.05. Fisher’s Least Significant Difference (LSD) test and Duncan test were used for the multiple comparison. Moreover, Principle Component Analysis (PCA) was performed to examine the similarities between different restoration patterns, and Redundancy analysis (RDA) was to explore the soil environmental parameters that significantly affecting the metabolic diversity of soil microbial communities. These analyses were performed in Vegan packages in R 3.2.4 (http://www.r-project.org).

## Results

### Soil characteristics

There was no significant difference in the SWC of the surface soil in dormant season between GL, FL, RL and PL patterns, while in growth season, the SWC of RL and PL were significantly higher than that of GL (*P* < 0.05) (Table 3). Overall, the SWC of GL was the lowest in the growing season, and FL obtained the highest value in the dormant season, which was about 4.6 times of the lowest. Soil pHs in GL and PL in dormant season were significantly lower than those in FL and RL, while in growing season, the pH was obviously higher in RL than in the other three patterns (*P* < 0.05). The contents of NH_4_^+^-N, NO_3_^−^-N and DOC were the highest in PL in the dormant season, whereas in the growth season, the highest values of NH_4_^+^-N and NO_3_^−^-N were in FL, and of DOC was in PL (*P* < 0.05).

**Table 3.** Effects of ecological restoration to soil physicochemical properties in different patterns.

different patterns
| Patterns | Seasons | SWC | pH | SOC | DOC | TN | NH<sub>4</sub><sup>+</sup>-N | NO<sub>3</sub><sup>-</sup>-N | TP | TK |
| --- | --- | --- | --- | --- | --- | --- | --- | --- | --- | --- |
|  |  | (%) |  | (g·kg<sup>-1</sup>) | (mg·kg<sup>-1</sup>) | (g·kg<sup>-1</sup>) | (mg·kg<sup>-1</sup>) | (mg·kg<sup>-1</sup>) | (g·kg<sup>-1</sup>) | (g·kg<sup>-1</sup>) |
| GL | DS | 28.28 ± | 6.42 ± | 6.59 ± | 19.99 ± | 0.66 ± | 2.05 ± | 0.35 ± | 0.05 ± | 4.16 ± |
|  |  | 0.73a | 0.10c | 1.09c | 7.55b | 0.10c | 0.41ab | 0.11bc | 0.00a | 0.49b |
|  | GS | 6.93 ± | 6.32 ± | 9.49 ± | 30.38 ± | 0.82 ± | 0.81 ± | 0.05 ± | 0.22 ± | 6.63 ± |
|  |  | 0.80B | 0.03B | 0.29B | 4.05AB | 0.04B | 0.14B | 0.01B | 0.02C | 2.74A |
| FL | DS | 31.69 ± | 7.14 ± | 23.92 ± | 35.68 ± | 1.86 ± | 2.02 ± | 0.60 ± | 0.05 ± | 7.67 ± |
|  |  | 3.49a | 0.26b | 1.52a | 2.31ab | 0.09a | 0.25ab | 0.18ab | 0.02a | 0.20a |
|  | GS | 11.39 ± | 6.21 ± | 23.81 ± | 32.31 ± | 1.75 ± | 2.07 ± | 1.04 ± | 0.67 ± | 5.11 ± |
|  |  | 0.95AB | 0.05B | 0.86A | 6.26AB | 0.05A | 0.14A | 0.26A | 0.04A | 0.37AB |
| RL | DS | 27.38 ± | 8.27 ± | 5.06 ± | 22.98 ± | 0.44 ± | 1.36 ± | 0.19 ± | 0.06 ± | 3.36 ± |
|  |  | 3.98a | 0.19a | 2.14c | 6.88ab | 0.12c | 0.13b | 0.01c | 0.01a | 0.74b |
|  | GS | 17.19 ± | 7.61 ± | 2.82 ± | 17.59 ± | 0.26 ± | 0.49 ± | 0.24 ± | 0.24 ± | 1.64 ± |
|  |  | 2.90A | 0.14A | 0.48C | 2.78B | 0.02C | 0.11B | 0.16B | 0.03C | 0.52B |
| PL | DS | 28.86 ± | 5.97 ± | 12.75 ± | 40.46 ± | 0.99 ± | 2.74 ± | 0.73 ± | 0.09 ± | 7.01 ± |
|  |  | 0.90a | 0.14c | 0.28b | 4.83a | 0.06b | 0.30a | 0.06a | 0.03a | 0.36a |
|  | GS | 13.43 ± | 6.24 ± | 9.16 ± | 36.43 ± | 0.91 ± | 1.81 ± | 0.52 ± | 0.39 ± | 5.48 ± |
|  |  | 1.56A | 0.30B | 0.66B | 6.17A | 0.05B | 0.48A | 0.07B | 0.01B | 0.24AB |

Mean values ± Stand.Error (n=3).
Small letters in the same column indicate the significant difference between restoration patterns
in the dormant season (DS), and capital letters indicate that in growing season (GS) ( $P < 0.05$ ).

The contents of SOC and TN were significantly higher in FL than in the other three patterns in both seasons (*P* < 0.05). The variation trend of SOC content during the dormant season was FL>PL>GL>RL, and that in growth season was FL>GL>PL>RL. For TN, the variation trend was FL>PL>GL>RL in both seasons. Furthermore, TP content did not change significantly in the dormant season, but showed significant differences in the growing season with the change trend of FL>PL>RL>GL. Generally among the four patterns, TP content in the growing season was higher than that in the dormant season. Besides, the variation range of TK content among the four patterns was 1.64-7.67 g·kg^−1^. Specifically, the TK contents were significantly higher in FL and PL than in GL and RL in the dormant season, while in the growing season, TK content in GL was significantly higher than those in RL and PL (*P* < 0.05).

### Microbial activity and physiological diversity

Carbon substrate utilization, assessed via Biolog method, showed that different restoration patterns modified the metabolic potential of the lakeside wetland soil microbial community, and this effect varied in different seasons. Overall, the AWCD values generally followed the same pattern with incubation time (leveling off after rapid increase). As shown in Fig 1, AWCD values of the four patterns showed a rapid growth trend at the initial stage of culture, with the fastest change rate at 72 h in both seasons. Specially, the AWCD value of RL pattern was always the highest at the detection points of each culture time, while the GL was the lowest.

**Fig 1.**
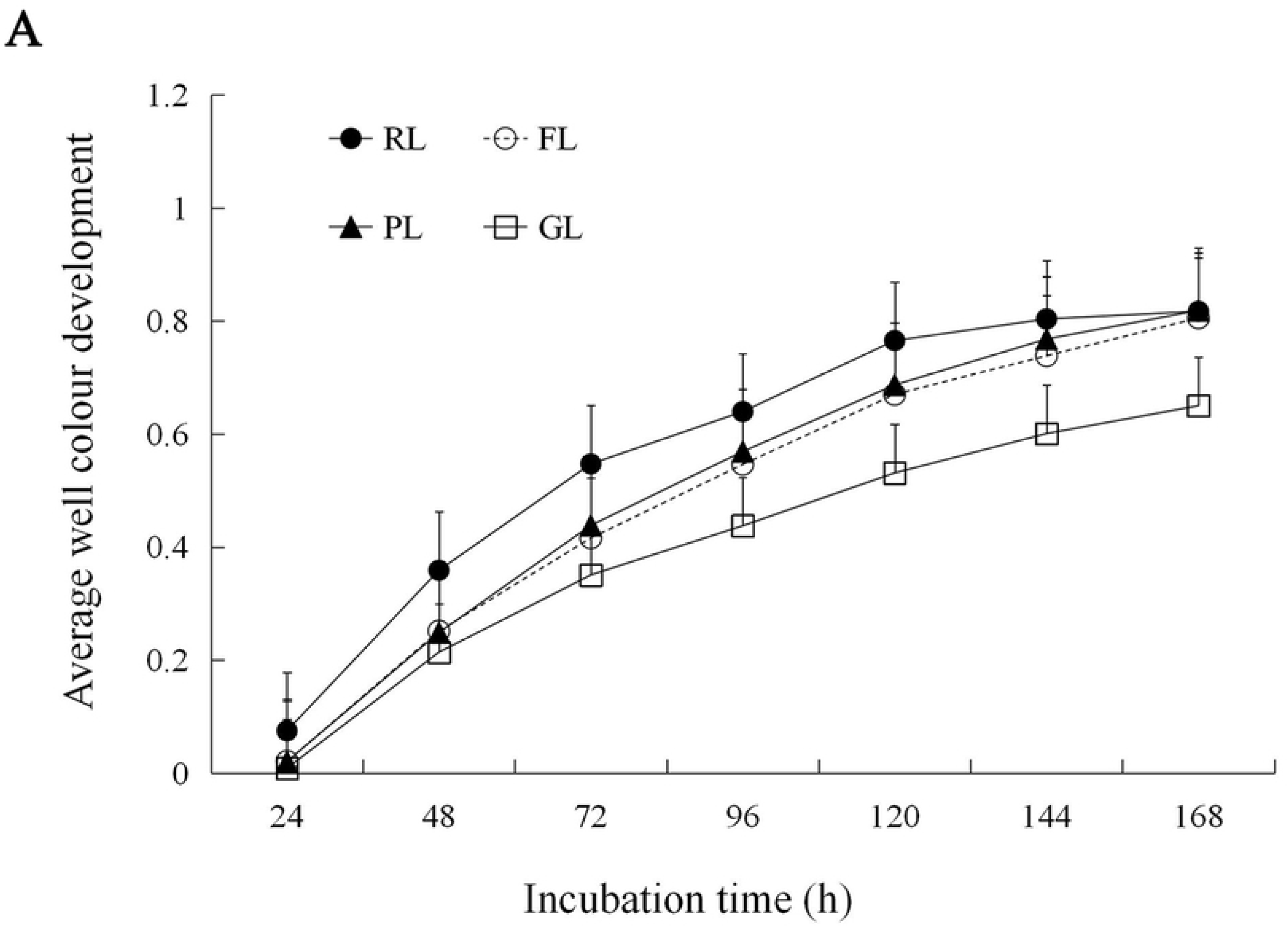
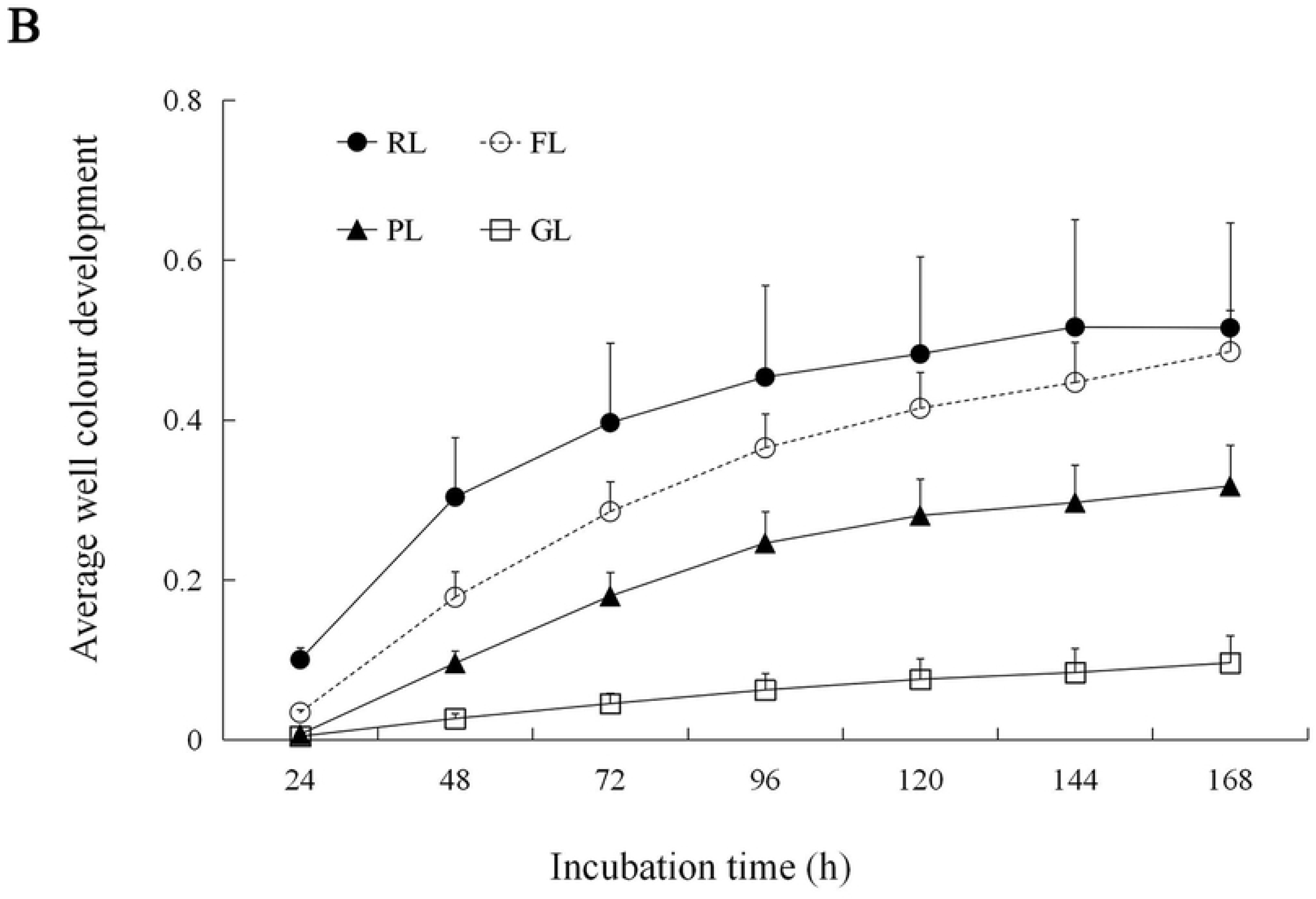
Average well color development (AWCD) of metabolized substrates in Biolog EcoPlates based on 168 h incubation (n=3). A-Dormant season, B-Growing season.

The functional diversity indices of soil microbial community was calculated based on AWCD values (Table 4). Results showed that in the dormant season, only the McIntosh index (*U*) obviously differed within the four patterns, which indicated that the *U* value in RL was significantly higher than that in GL, but not significantly differed with FL or PL (*P* < 0.05). During the growing season, however, RL pattern had higher Shannon-Wiener index (*H*′) compared to the other patterns (*P* < 0.05), and it ranked in the following order: RL>FL>PL>GL. Besides, no significant difference on McIntosh index and Simpson index was observed in this time.

**Table 4.**
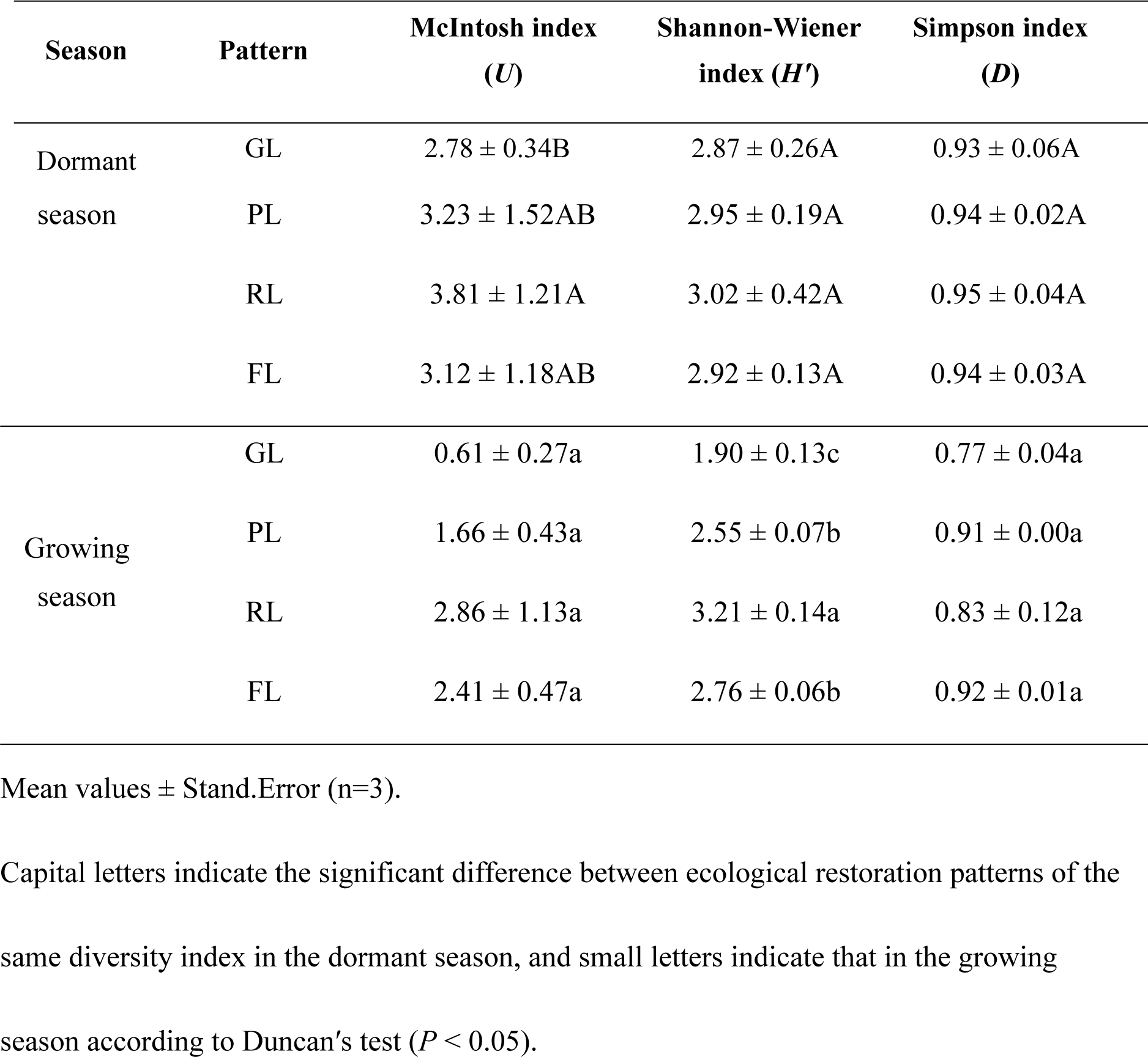
Diversity indices based on the carbon-source utilization model for the soil samples from GL, PL, RL and FL.

| Season | Pattern | McIntosh index<br>($U$) | Shannon-Wiener<br>index ($H'$) | Simpson index<br>($D$) |
| --- | --- | --- | --- | --- |
| Dormant | GL | 2.78 ± 0.34B | 2.87 ± 0.26A | 0.93 ± 0.06A |
| season | PL | 3.23 ± 1.52AB | 2.95 ± 0.19A | 0.94 ± 0.02A |
|  | RL | 3.81 ± 1.21A | 3.02 ± 0.42A | 0.95 ± 0.04A |
|  | FL | 3.12 ± 1.18AB | 2.92 ± 0.13A | 0.94 ± 0.03A |
| Growing<br>season | GL | 0.61 ± 0.27a | 1.90 ± 0.13c | 0.77 ± 0.04a |
|  | PL | 1.66 ± 0.43a | 2.55 ± 0.07b | 0.91 ± 0.00a |
|  | RL | 2.86 ± 1.13a | 3.21 ± 0.14a | 0.83 ± 0.12a |
|  | FL | 2.41 ± 0.47a | 2.76 ± 0.06b | 0.92 ± 0.01a |

Mean values ± Stand.Error (n=3).
Capital letters indicate the significant difference between ecological restoration patterns of the same diversity index in the dormant season, and small letters indicate that in the growing season according to Duncan's test ( $P < 0.05$ ).

The EcoPlate carbon sources were divided into six biochemical categories, and the results from the 72-h incubation showed that in the dormant season, the relative utilization rate of carboxylic acids in RL was at a lower level, and that of amines was high. While the opposite trend was observed in GL, that is, carboxylic acids were most widely used, whereas the utilization of amines was less. Furthermore, soil microbial communities in FL and PL patterns had an average utilization efficiency for different carbon substrates (Fig 2A). During the growing season, the relative utilization rate of phenolic acids in GL was observed to be far more less than that in other three patterns, while the rate on carboxylic acids was significantly higher compared with RL, FL and PL (Fig 2B).

**Fig 2.**
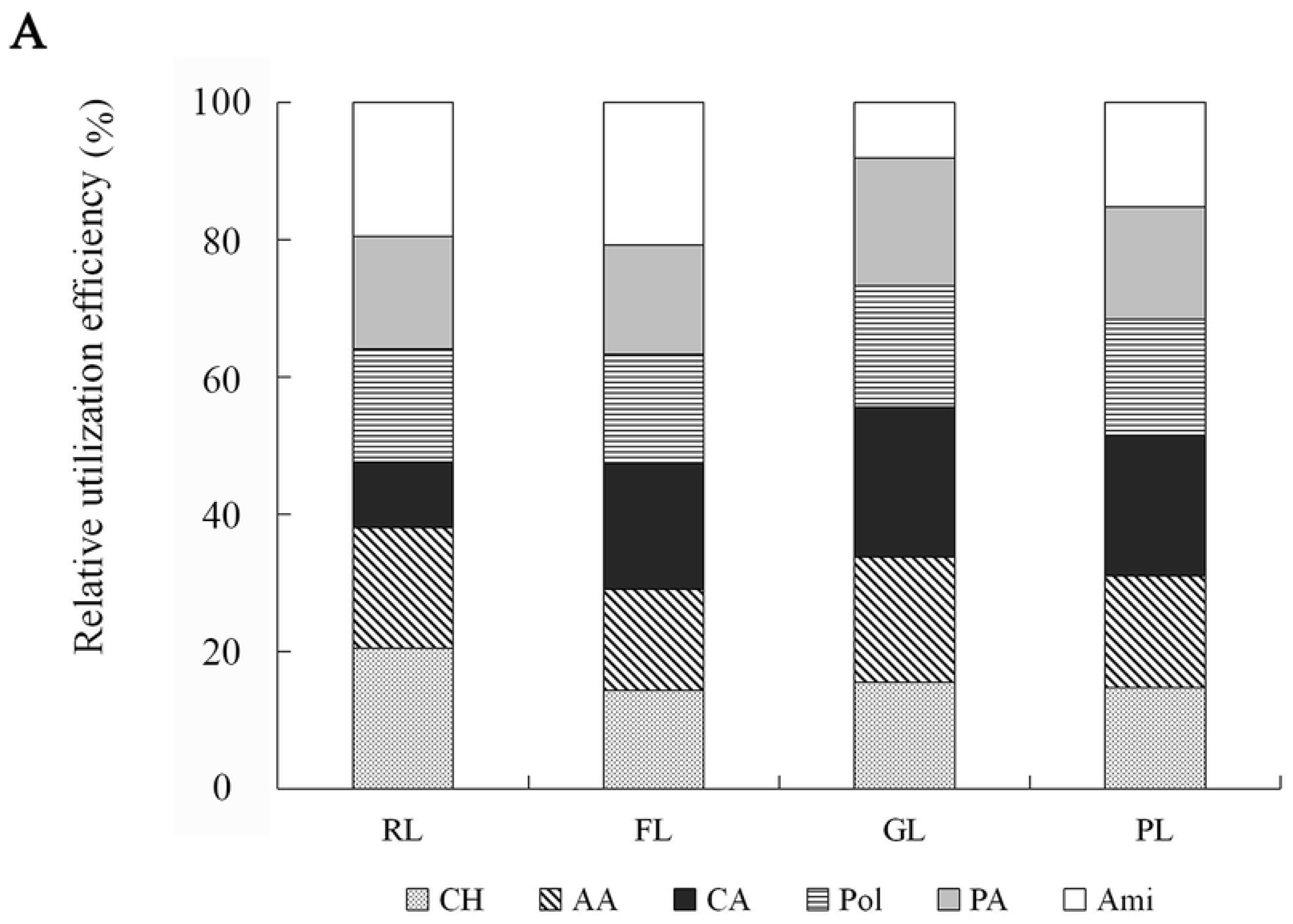
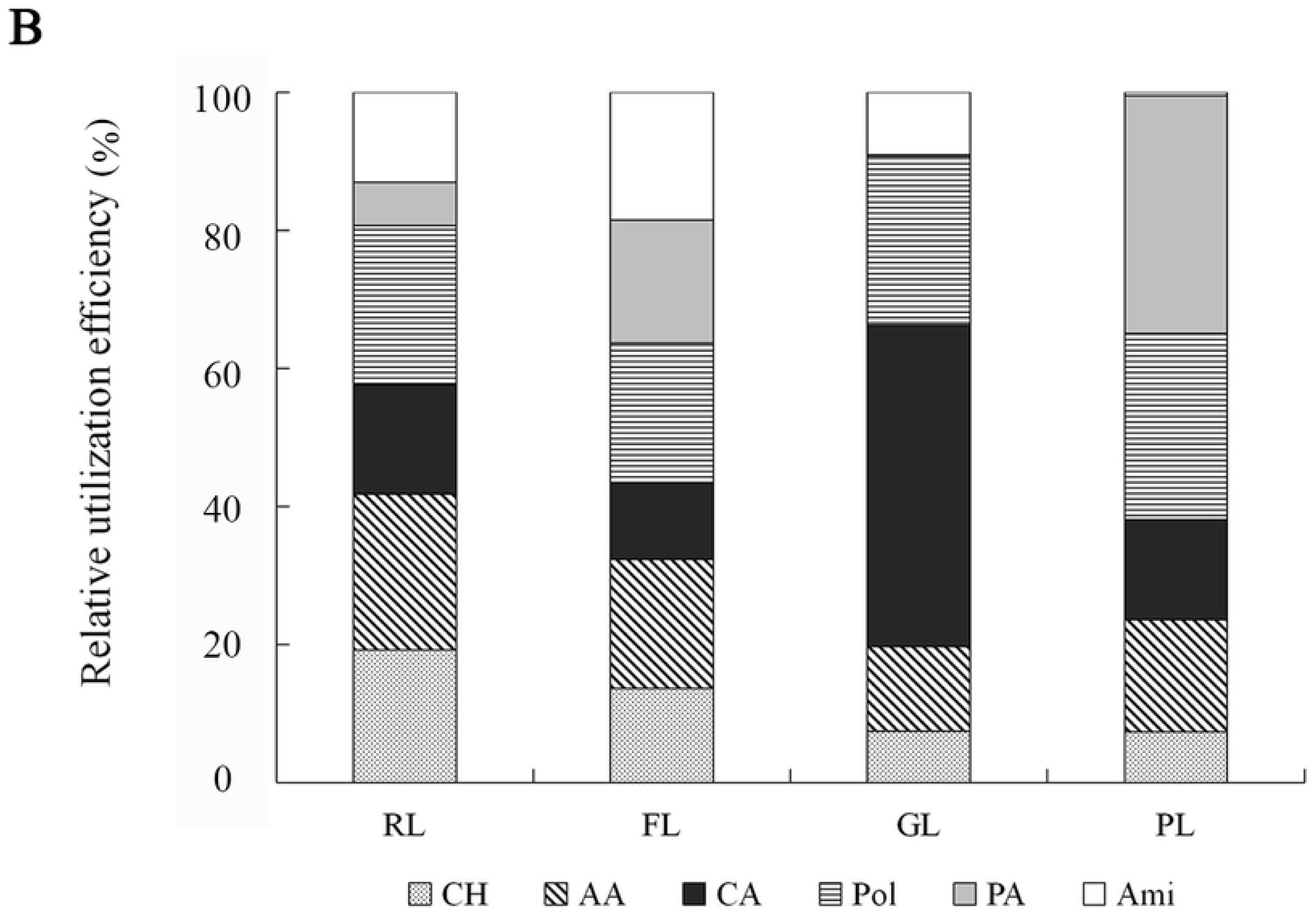
The relative utilization efficiency of carbon sources in soil microbial community. (a) CH-Carbohydrates, AA-Amino acids, CA-Carboxylic acids, PA-Phenolic acids, Pol-Polymers, Ami-Amines. (b) A-Dormant season, B-Growing season.

### PCA analysis of metabolic characteristics of soil microbial communities

In order to determine the level of sites differentiation, the PCA ordination plots were performed though CLPPs. In the dormant season, the cumulative contribution rate of the first two axes was up to 77.49%, with PCA1 and PCA2 axes reaching 63.32% and 14.17%, respectively (Fig 3). Among the four ecological restoration patterns, GL and RL were distributed in a concentrated way, and the samples in GL, FL and PL grouped together which were distinct from samples in RL, indicating that the soil microbial communities in GL, FL and PL had similar carbon-source utilization patterns and were significantly different from those in RL.

**Fig 3.**
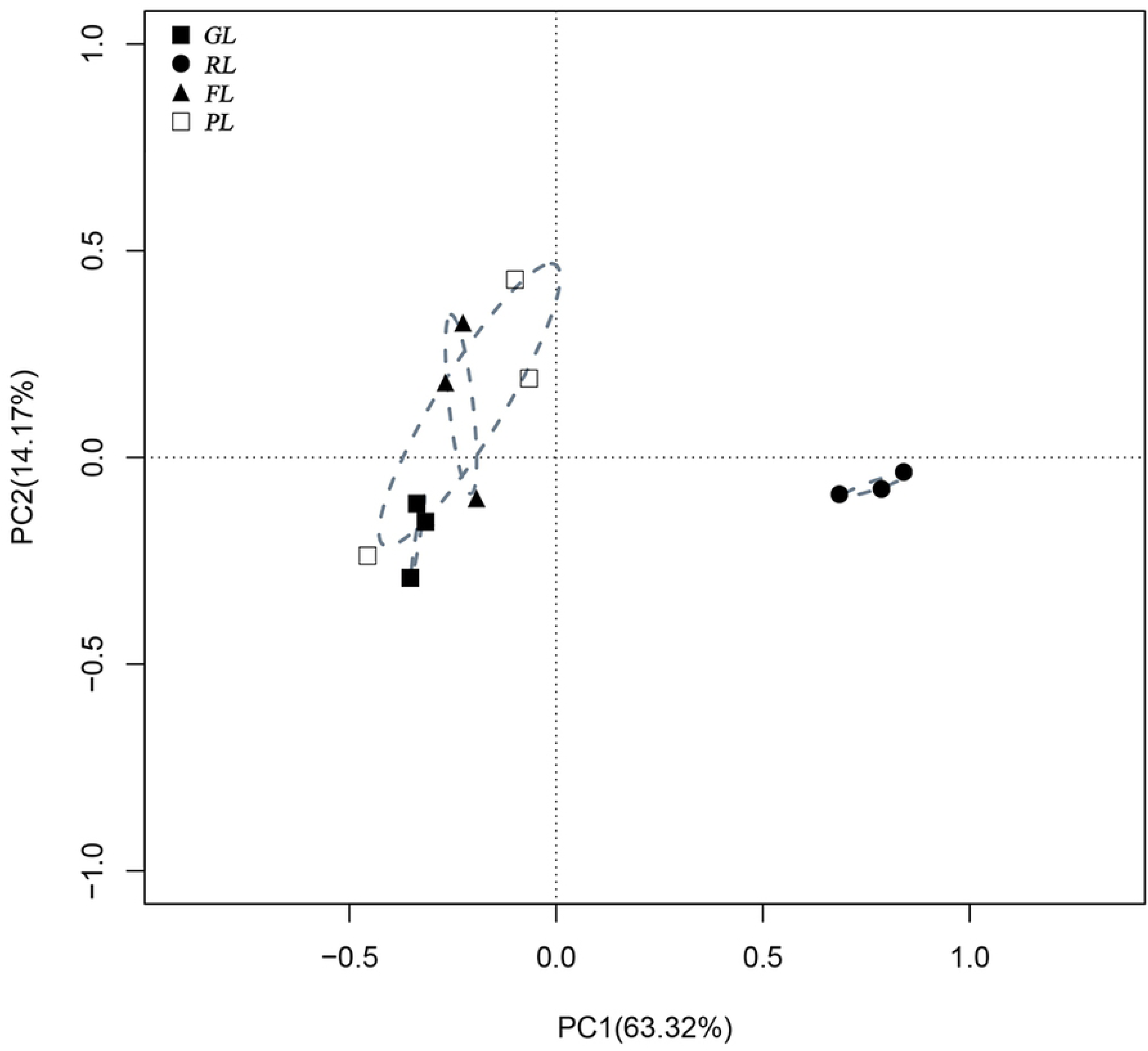
PCA analysis of soil microbial carbon-source utilization in dormant season. The dotted circles represent the 95% confidence interval.

The load values of 31 carbon sources on the two principal components were shown in Table 5. It can be seen that there were 4 types of carbon sources significantly correlated with PC1, and 10 types with PC2 (*P* < 0.05). Consequently, carbohydrates were the main carbon sources that distinguished the soil microbial metabolic characteristics from differential restoration patterns in dormant season.

**Table 5.** Correlation coefficients of 31 carbon sources with PC1 and PC2 in different seasons.

| Category | Carbon sources (Code) | Dormant season |  | Growing season |  |
| --- | --- | --- | --- | --- | --- |
|  |  | PC1 | PC2 | PC1 | PC2 |
| Carbohydrates | β-Methyl-D-Glucoside (CH1) | 0.6515 | -0.0845 | 0.2163 | **0.0066** |
| | D-Galactonic Acid-γ-Lactone (CH2) | 0.2391 | **-0.0042** | 0.2572 | -0.0522 |
|  | D-Xylose (CH3) | 0.1023 | **-0.0026** | **0.0213** | 0.3812 |
|  | i-Erythritol (CH4) | -0.1165 | 0.1768 | 0.0974 | **0.0172** |
|  | D-Mannitol (CH5) | 0.1555 | 0.3133 | 0.7302 | -0.0844 |
|  | N-Acetyl-D-Glucosamine (CH6) | 0.5176 | **-0.0098** | 0.3698 | 0.3988 |
|  | D-Cellobiose (CH7) | 0.3415 | 0.1551 | 0.5898 | -0.1076 |
| | α-D-Glucose-1-Phosphate (CH8) | 0.1052 | **-0.0170** | 0.3914 | 0.3516 |
| | α-D-Lactose (CH9) | 0.1495 | **0.0263** | 0.0977 | 0.0765 |
| | D,L-α-Glycerol Phosphate (CH10) | **0.0113** | **0.0127** | 0.0686 | 0.2336 |
| Amino acids | L-Arginine (AA1) | 0.4320 | **0.0348** | 0.4982 | -0.2069 |
|  | L-Asparagine (AA2) | 0.2791 | 0.1894 | 0.5797 | -0.1466 |
|  | L-Phenylalanine (AA3) | -0.1487 | **0.0207** | 0.0779 | 0.3761 |
|  | L-Serine (AA4) | 0.7437 | -0.0739 | 0.3912 | **0.0070** |
|  | L-Threonine (AA5) | -0.1504 | 0.1903 | 0.3222 | 0.2502 |
|  | Glycyl-L-Glutamie Acid (AA6) | **-0.0174** | **0.0261** | **0.0138** | **0.0178** |
| Carboxylic acids | Pyruvic Acid Methyl Ester (CA1) | **0.0334** | 0.2546 | 0.4467 | -0.1693 |
|  | D-Galacturonic Acid (CA2) | -0.1068 | 0.1008 | 0.4313 | **-0.0132** |
| | γ-Hydroxybutyric Acid (CA3) | **0.0319** | 0.2510 | **0.0285** | 0.1643 |
|  | D-Glucosaminic Acid (CA4) | -0.1013 | 0.1908 | -0.0578 | 0.3297 |
|  | Itaconic Acid (CA5) | 0.1867 | 0.0872 | 0.1654 | **-0.0193** |
| | α-Ketobutyric Acid (CA6) | -0.1108 | 0.0829 | **0.0381** | 0.1511 |
|  | D-Malic Acid (CA7) | -0.6527 | 0.1955 | 0.1709 | **0.0277** |
| Polymers | Tween 40 (Pol1) | 0.1734 | 0.1440 | 0.2887 | 0.4086 |
|  | Tween 80 (Pol2) | 0.1373 | 0.0731 | 0.3898 | **0.0338** |
| | α-Cyclodextrin (Pol3) | 0.1130 | 0.0731 | **0.0268** | 0.2874 |
|  | Glycogen (Pol4) | 0.0906 | **0.0028** | 0.4082 | -0.0541 |
| Phenolic acids | 2-Hydroxy Benzoic Acid (PA1) | -0.0548 | 0.0911 | -0.1231 | 0.5939 |
|  | 4-Hydroxy Benzoic Acid (PA1) | 0.2847 | 0.1370 | 0.19329 | -0.0766 |
| Amines | Phenylethyl-amine (Ami1) | 0.1725 | 0.1239 | 0.1489 | **-0.0017** |
|  | Putrescine (Ami1) | 0.2075 | 0.1563 | 0.3163 | -0.1461 |

Bold values indicates significant difference at the level of 0.05.

Considering the microbial communities in the growing season, PCA analysis showed that the contribution rate of PCA1 and PCA2 axes reached 52.19% and 26.27%, respectively (Fig 4). Among the four patterns, samples in GL and PL distributed in a concentrated way, whereas samples in FL and RL were relatively discrete and crossed each other. Besides, by calculating the loading values, we found 5 types of carbon sources highly correlating with PC1, and 9 types with PC2 (*P* < 0.05, Table 5). Combined with PC1 and PC2, carboxylic acids played a leading role in the metabolic characteristics of soil microbial communities among different restoration patterns in the growing season.

**Fig 4.**
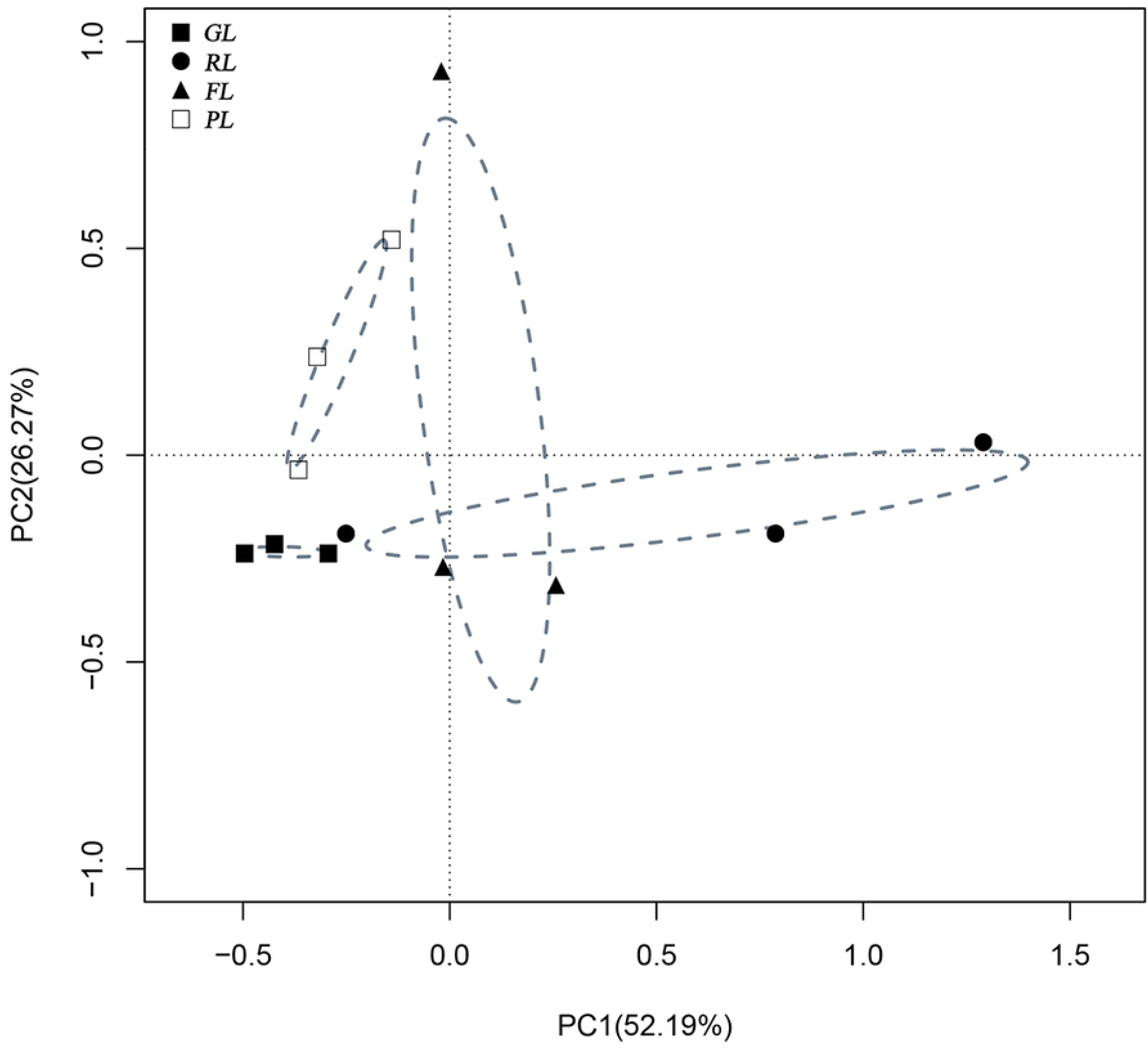
Principal component analysis of soil microbial carbon-source utilization in the growing season. The dotted circles represent the 95% confidence interval.

### RDA analysis on effect of soil environmental factors on microbial functional diversity

In the dormant season, the first two RDA axes explained 70.68% of the carbon-source utilization under the constraint ordination (Figure 5). The solid lines of soil pH and TK were shown to be longer than other parameters, which indicated their serious impacts on the microbial metabolism. Monte Carlo permutation tests further verified these significant effects of pH and TK, with correlation coefficient r^2^ of 0.545 and 0.573, respectively (*P* < 0.05, Table 6). On the other hand, the dotted arrows showed that the carbon-source utilization in RL pattern were obviously different from PL, GL and FL that distributing in the positive axis of RDA1. It was consistent with the PCA results. The permutation tests further demonstrated the significant effects of different restoration patterns on functional diversity of soil microbial communities (r^2^ = 0.754, *P* < 0.01, Table 6).

**Fig 5.**
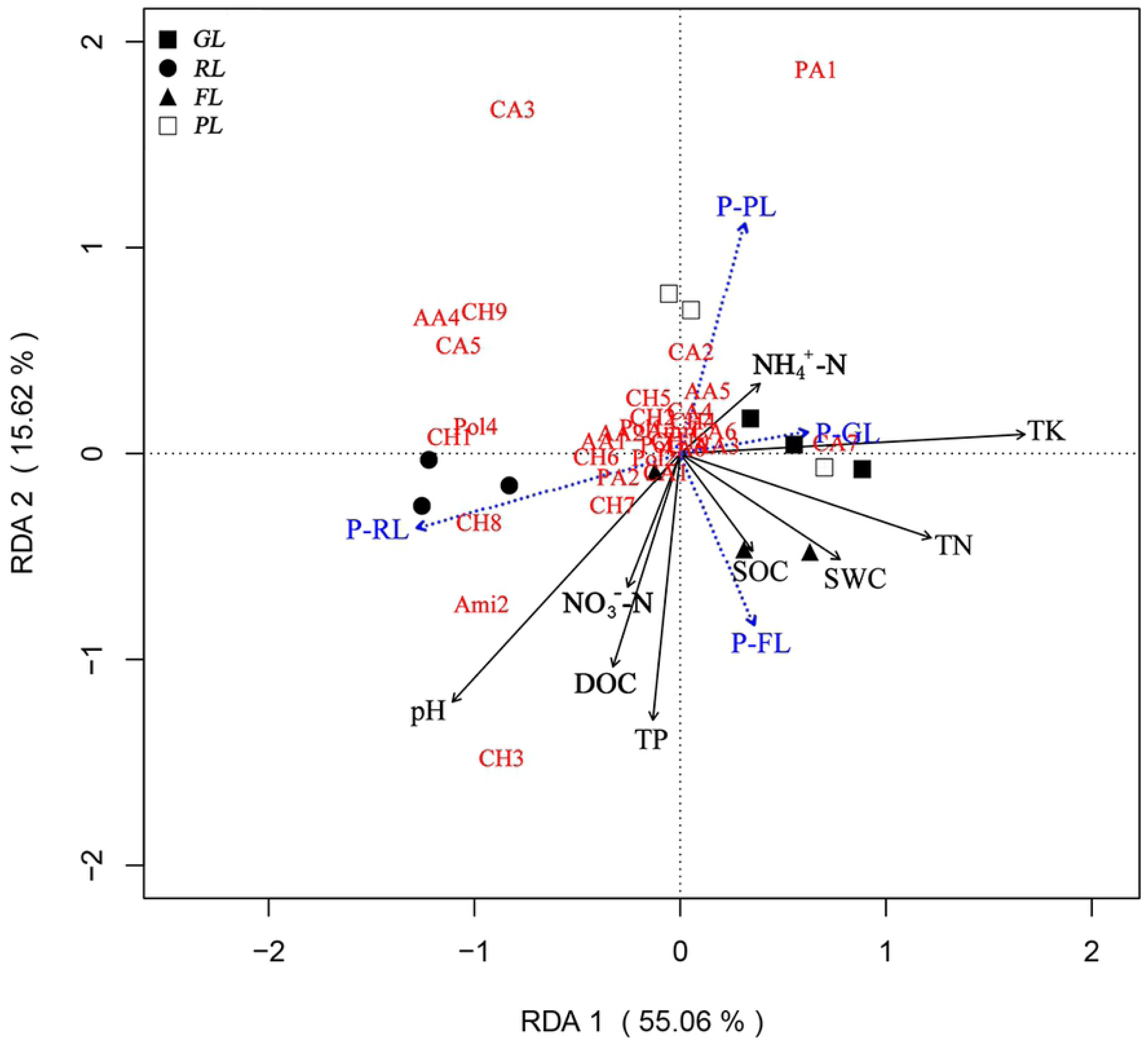
Redundancy analysis between the soil properties and microbial functional diversity characteristics in dormant season. Carbon-source code is the same as in Table 5.

**Table 6.**
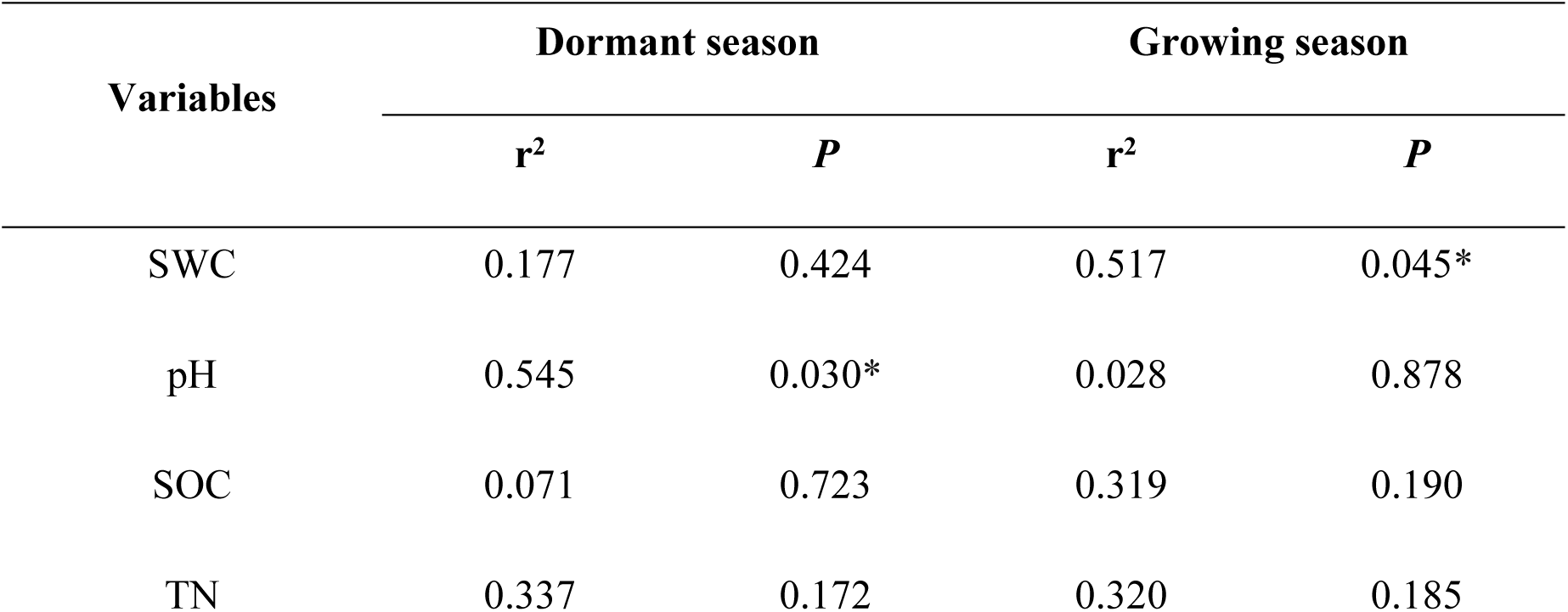
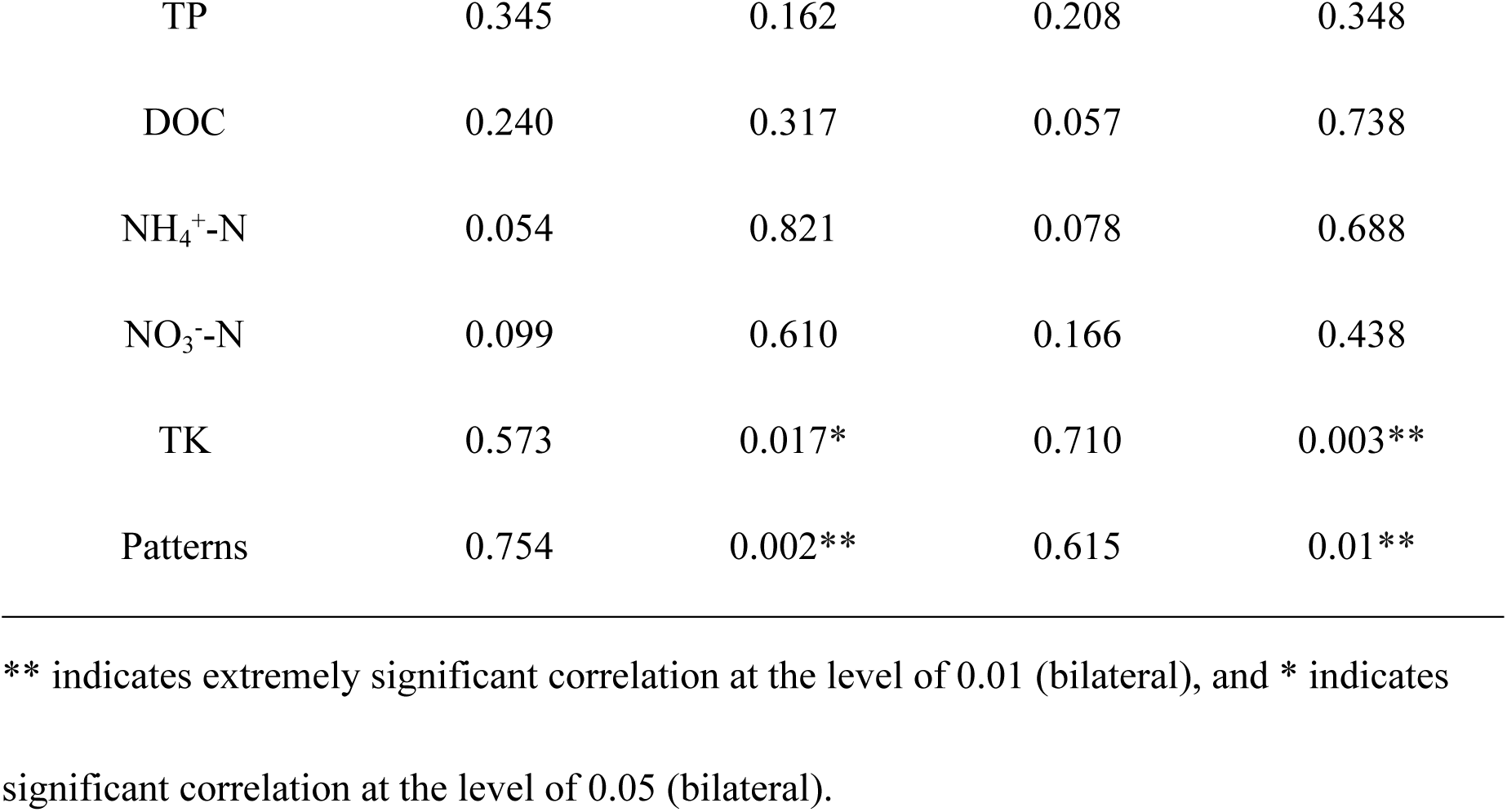
Monte Carlo permutation tests of the effect of environmental variables on microbial carbon-source utilization in dormant and growing seasons.

| Variables | Dormant season |  | Growing season |  |
| --- | --- | --- | --- | --- |
| | $r^2$ | $P$ | $r^2$ | $P$ |
| SWC | 0.177 | 0.424 | 0.517 | 0.045* |
| pH | 0.545 | 0.030* | 0.028 | 0.878 |
| SOC | 0.071 | 0.723 | 0.319 | 0.190 |
| TN | 0.337 | 0.172 | 0.320 | 0.185 |
| TP | 0.345 | 0.162 | 0.208 | 0.348 |
| DOC | 0.240 | 0.317 | 0.057 | 0.738 |
| NH<sub>4</sub><sup>+</sup>-N | 0.054 | 0.821 | 0.078 | 0.688 |
| NO<sub>3</sub><sup>-</sup>-N | 0.099 | 0.610 | 0.166 | 0.438 |
| TK | 0.573 | 0.017* | 0.710 | 0.003** |
| Patterns | 0.754 | 0.002** | 0.615 | 0.01** |

\*\* indicates extremely significant correlation at the level of 0.01 (bilateral), and \* indicates significant correlation at the level of 0.05 (bilateral).

Furthermore, the plot also showed that various types of carbon-source utilized by microorganisms in different patterns were quite similar, and concentrated in the center of the axis (Fig 5). Based on the distribution distance between sites and carbon sources, we observed that the microbial communities in RL used more carbohydrates, GL sites preferred carboxylic acids, while PL and FL took the average utilization of different carbon substrates because of the similar distance from most sources concentrating in the center point.

Based on the PCA and RDA analysis, the specific correlations between soil pH, TK and 31 types of carbon-sources were tested through Pearson’s correlation analysis (Table A1). The results showed that in the dormancy season, there was a extremely significant negative correlation between soil pH and L-Threonine (AA5, r^2^ = −0.714, *P* < 0.01), whereas soil TK had negative correlations with β-Methyl-D-Glucoside (CH1), D-Galactonic Acid-γ-Lactone (CH2) and L-Arginine (AA1) (*P* < 0.01). Overall, there were 6 in 10 kinds of carbohydrates and 5 in 6 of amino acids significantly correlated with TK content (*P* < 0.05).

In the growing season, the first two axes (RDA1 and RDA2) constructed the RDA plot, which could explain 48.85% and 19.86% of the total variation, respectively (Figure 6). The length of the solid arrow lines displayed that soil SWC and TK had significant effect on microbial carbon-source utilization, with r^2^ calculated in permutation test of 0.517 and 0.710, respectively (*P* < 0.05). The dotted lines showed the obvious differentiation between GL, PL and RL, whereas there were similarities of the metabolic diversity in RL and FL. Moreover, the distribution of the carbon-sources in RDA ordination plot displayed that the substrates utilized by microorganisms in different patterns were relatively discrete, and the sources that may be used by certain communities were largely scattered around the positive half of the RDA1 axis (Fig 6). Overall, amino acids, polymers and phenolic acids were increasingly utilized in RL, PL and FL patterns, which were different from the metabolic characteristics in GL.

**Fig 6.**
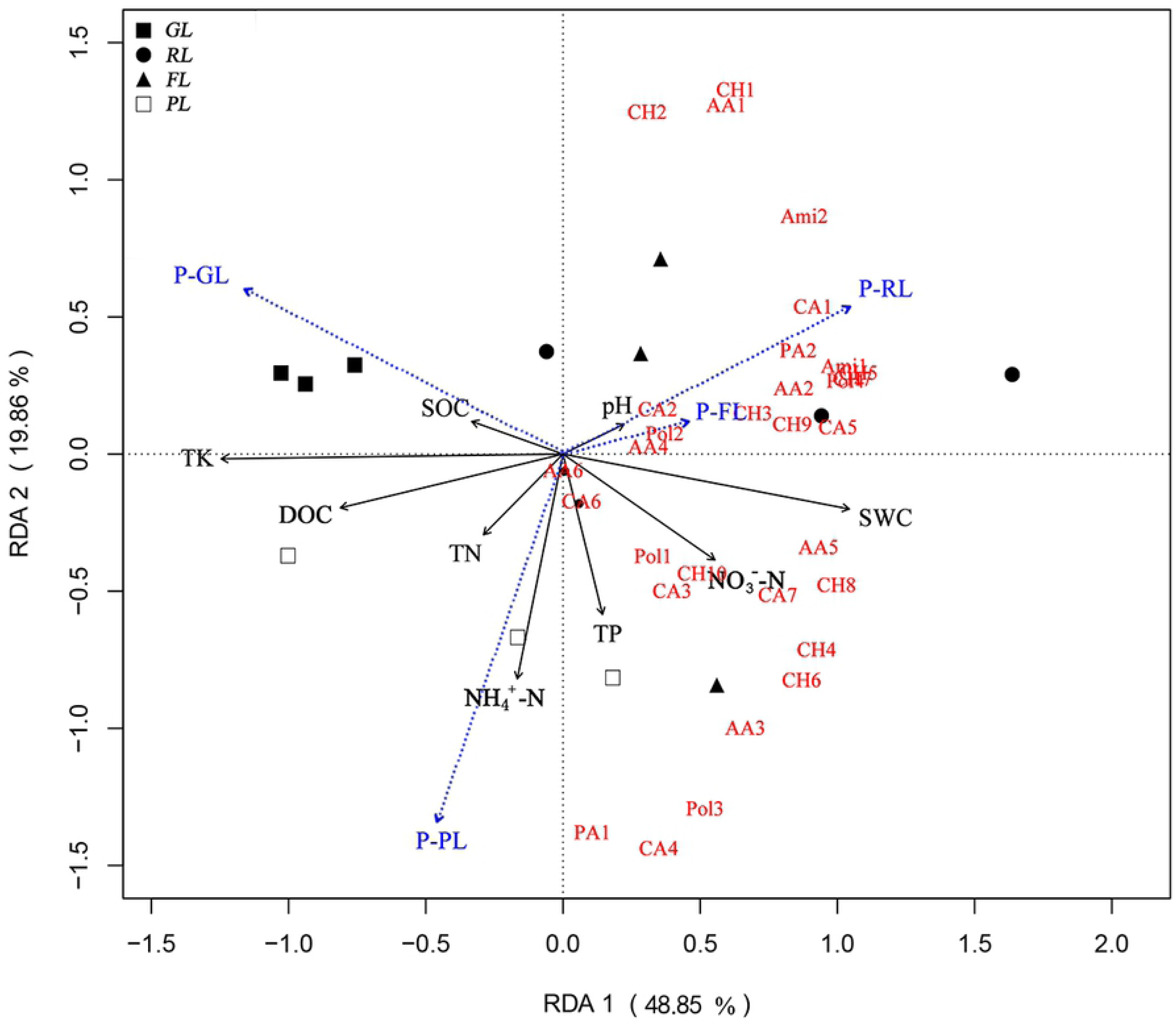
Redundancy analysis (RDA) between the soil properties and microbial functional diversity characteristics in growing season. Carbon-source code is the same as in Table 5.

Pearson’s correlation analysis indicated that in the growing season, SWC positively affected the utilization of several carbon sources, especially for i-Erythritol (CH4), D-Mannitol (CH5), N-Acetyl-D-Glucosamine (CH6) and D-Cellobiose (CH7) in carbohydrates (*P* < 0.01), and Pyruvic Acid Methyl Ester (CA1), Itaconic Acid (CA5) and D-Malic Acid (CA7) in carboxylic acids (*P* < 0.05). However, the correlations between soil TK and carbon sources were just the opposite, for the significant negative relations with i-Erythritol (CH4), D-Mannitol (CH5), N-Acetyl-D-Glucosamine (CH6), D-Cellobiose (CH7) and α-D-Glucose-1-Phosphate (CH8) in carbohydrates (*P* < 0.01), and D-Galacturonic Acid (CA2), Itaconic Acid (CA5) and D-Malic Acid (CA7) in carboxylic acids (*P* < 0.05) (Table in S1 Table).

## Discussion

In the present study, the Biolog EcoPlates™ method was used to monitor the soil microbial functional diversity of Chaohu lakeside wetland. The AWCD values recorded the overall ability of microbial communities to utilize 31 single carbon-sources within 168 h [34], showing that the carbon-source utilization rate increased significantly after 24 hours of incubation, and their utilization capacity was obviously different among four patterns. Overall the microbial metabolic activity was lower in GL pattern than in other three patterns, and the relatively lower diversity index also demonstrated the poorer microbial populations in GL. As the derelict land around the lake, GL sites are covered by only weed, without any trees and shrubs due to the heavy clay and hardening. Previous research has reported that the abandoned fields are often characterized by less diverse plant communities [35, 36]. Besides, soil C and N fractions are derived from the decomposition of litter [37], the release of root exudates, and rhizodeposition [38]. Lack of aboveground cover could decrease the quantity and quality of litter and roots, thus lowering the soil nutrient content and cycling, which may be negative for the soil microbial diversity.

In our study, soil microorganisms were observed to have various preferences on the carbon-source utilization in different seasons, especially for carbohydrates in dormant season and carboxylic acids in growing season. Carbohydrates, which belong to the soil active organic carbon pool [39], provide energy and substrates for microorganisms, and are significantly related to microbial activity [40, 41]. Thereby many aerobic and facultative heterotrophs choose carbohydrates as their main sources to realize their functions, the process of which involves the oxidation of simple or complex carbohydrates [42–45]. Moreover, carboxylic acids are the important part of the organic acid that can affect the release of heavy metal ions in soil and improve their activities [46]. Lin et al. [47] found that the increase of heavy metals and their availability in soil enhanced the metabolic utilization of carboxylic acids. In summer, water eutrophication may lead to more nutrients enriching into the wetland soil, which may stimulate the microorganisms to use more carboxylic acids to adsorb and degrade heavy metals. This is consistent with Li et al. [48] who reported that carboxylic acids and carbohydrates were the sensitive carbon-sources affecting the metabolic function of microbial communities in the Songjiang wetland under the effect of four kinds of disturbances.

Interestingly, when taking the environmental factors into consideration, the situation of functional metabolism became much more complicated in various restoration patterns from RDA results. In dormant season, the carbon sources like carbohydrates and carboxylic acids were used widely, and the microbial communities in four patterns showed a similar way in selecting substrates. However, the microbial communities showed more versatile substrate utilization patterns in growing season than in dormant season. For example, RL increased the substrate utilization of amino acids, whereas microorganisms in PL and FL used more phenolic acids and polymers. Consequently, not only the easily degraded compounds but also the complex ones were consumed to satisfy the metabolic requirement of soil microorganisms. Adam et al. [12] also reported that in spring and summer, the frequency of nitrogen-rich carbon sources (amino acids) [45] used by microbial communities was higher than that of carbohydrates and carboxylic acids. In addition to utilizing carbon, these microbial communities also use amino acids as nitrogen source and combine with ammonia side chains. Ammonium is then made into organic molecules, such as amino acids and proteins [49]. Moreover, phenolic acids are a major class of phenolic compounds made by plants [50], and denaturing gradient gel electrophoresis (DGGE) results obtained by Zhou and Wu [51, 52] have proved that the phenolic acid supplementations can significantly alter the soil bacterial community composition. Besides, polymers are complex carbon substrates, and previous research has shown that when the growth of soil bacteria was inhibited due to lack of required nutrients (e.g., nitrogen or phosphate) in the presence of excess carbon, the glycogen may accumulate in bacteria [53]. In our study, the high use of these groups (amino acids, phenolic acids and polymers) in RL, PL and FL patterns might be related to the availability of various substrates in growing season. After vegetation restoration, soil microbial populations and diversity increased significantly compared to the shoaly abandoned land, which stimulate their competition for utilizing different kinds of substrate, thus enhance the complex carbon-source degradation under the suitable hydrothermal condition to satisfy their multiple functional needs.

Furthermore, CLPPs seem to be specific for land use, soil management and soil texture [13]. Our research showed that soil TK contents was the dominant factor affecting the microbial functional diversity in both dormant and growing seasons. After nitrogen and phosphorus, potassium (K) is one of the major nutrients required by vegetation, and it plays a major role in the activation of several metabolic processes including protein synthesis, photosynthesis, enzyme activation, etc [54]. Specially, K acts as a key to activate enzymes to metabolize carbohydrates for the synthesis of amino acids and proteins [55], and previous studies reported diverse groups of soil microorganisms took an active part in solubilizing the fixed forms of K into available forms which plants are easy to absorb [56, 57]. This may explain the significant relationship between TK and substrates of carbohydrates and amino acids used by soil microbial communities observed in our study. The utilization ability of communities to different carbon-sources, however, depends to a large extent on the types and inherent properties of microorganisms, as Barra Caracciolo et al. [58] observed the shifts of soil microbial composition determined their functional diversity, and finally had different impacts on the ecosystem processes. The EcoPlates™ method can be used to describe the utilization of microbial communities to a single type of carbon-sources, whereas the complex edaphic environment could not be simulated. Therefore, future research on microbial compositional and structural information is needed to help to understand the mechanism about the metabolic shifts of microbial communities during the ecological restoration in Chaohu lakeside wetland.

## Conclusions

This study showed that Chaohu lakeside wetland soils with different biogeochemical properties had microbial communities that exhibit distinct catabolic responses to a range of carbon-sources. The AWCD values and diversity indices indicated soils in RL, PL and FL patterns had the higher overall metabolic activity of different substrates, whereas the microbial communities in GL had the relative lower ability. Seasonal variation of functional diversity was observed between dormant and growing seasons, and the CLPPs results clearly distinguished the metabolic characters in natural and artificial restoration patterns from that in the abandoned shoaly land by their capacity to utilize more complex carbon sources, such as amino acids, phenolic acids and polymers, which may be linked to differences in the soil microbial composition, vegetation types, soil physicochemical properties and hydrothermal condition, etc. All the soil parameters and Biolog data demonstrate the positive effect of ecological restoration on increasing the microbial activity and functional diversity. This study also displayed a close linkage between physicochemical properties (TK, pH and SWC) of lakeside wetland soil and the associated microbial functional activities, which may provide basic information regarding the environmental protection and management of Chaohu wetland.

## Author Contributions

Conceptualization: ZT.

Data curation: ZT.

Formal analysis: WF HW.

Investigation: HW XC.

Methodology: ZT WF.

Supervision: XX.

Writing – original draft: ZT.

Writing – review & editing: XX.

## Funding

This work was funded by the National Natural Science Foundation of China (NSFC, No. 31770672 and 31370626, received by XX), and the Graduate Innovation Foundation of Anhui Agricultural University (2018yjs-18, received by ZT).

## Acknowledgments

We would like to thank Xinlong Huang, Liang Wang, Peixi Li and Mengyao Sun for assistance with the fieldwork.

## Data Availability Statement

All relevant data are within the paper and its Supporting Information files.

## Conflicts of Interest

The authors have declared that no competing interests exist.

## Supporting information

**S1 Table. Correlation coefficients of the microbial carbon-source utilization with the soil environmental parameters in dormant and growing seasons.** ** indicates extremely significant correlation at the level of 0.01 (bilateral), and * indicates significant correlation at the level of 0.05 (bilateral). The carbon-source code is the same as in Table 3.

## References

1. Bernal B, Mitsch WJ. A comparison of soil carbon pools and profiles in wetlands in Costa Rica and Ohio. Ecol Eng. 2008;34(4):311–23.

2. Jiang M, Lv X, Yang Q, Tong S. Iron biogeochenical cycle and its environmental effect in wetlands (in Chinese). Acta Pedologica Sinica. 2006;43(3):493–9.

3. Ma Z, Cai Y, Li B, Chen J. Managing Wetland Habitats for Waterbirds: An International Perspective. Wetlands. 2010;30(1):15–27.

4. Withey P, Kooten GCV. The effect of climate change on optimal wetlands and waterfowl management in Western Canada. Ecol Econ. 2011;70(4):798–805.

5. Niemuth ND, Fleming KK, Reynolds RE. Waterfowl conservation in the US Prairie Pothole Region: confronting the complexities of climate change. PLoS One. 2014;9(6):e100034.

6. Mitsch WJ, Wu X, Nairn RW, Weihe PE, Wang N, Deal R, et al. Creating and restoring wetlands. Bioscience. 1998;48(12):1019–30.

7. Hou C. Effects of hydrological changes on soil carbon sequestration of marsh in Sanjiang Plain (in Chinese). Beijing: Chinese Academy of Sciences; 2012.

8. Zou Y, Ming J, Yu X, Lu X, David JL, Wu H. Distribution and biological cycle of iron in freshwater peatlands of Sanjiang Plain, Northeast China. Geoderma. 2011;164(3):238–48.

9. Zou YC, Xiao-Fei YU, Huo LL, Xian-Guo L, Jiang M. Waterborne Iron Migration by Groundwater Irrigation Pumping in a Typical Irrigation District of Sanjiang Plain (in Chinese). Environmental Science 2012;33(4):1209–15.

10. Selivanovskayaab SY. Ecotoxicological assessment of soil using the Bacillus pumilus contact test. Eur J Soil Biol. 2011;47(2):165–8.

11. Galitskaya P, Biktasheva L, Saveliev A, Ratering S, Schnell S, Selivanovskaya S. Response of soil microorganisms to radioactive oil waste: results from a leaching experiment. Biogeosciences,12,12(2015-06-16). 2015;12(2):1753–89.

12. Oest A, Alsaffar A, Fenner M, Azzopardi D, Tiquia-Arashiro SM. Patterns of Change in Metabolic Capabilities of Sediment Microbial Communities in River and Lake Ecosystems. Int J Microbiol. 2018;2018:1–15.

13. Rutgers M, Wouterse M, Drost SM, Breure AM, Mulder C, Stone D, et al. Monitoring soil bacteria with community-level physiological profiles using Biolog™ ECO-plates in the Netherlands and Europe. Appl Soil Ecol. 2016;97:23–35.

14. Feigl V, Ujaczki E, Vaszita E, Molnar M. Influence of red mud on soil microbial communities: Application and comprehensive evaluation of the Biolog EcoPlate approach as a tool in soil microbiological studies. Sci Total Environ. 2017;595:903–11.

15. Jiang L, Han G, Lan Y, Liu S, Gao J, Xu Y, et al. Corn cob biochar increases soil culturable bacterial abundance without enhancing their capacities in utilizing carbon sources in Biolog Eco-plates. Journal of Integrative Agriculture. 2017;16(3):713–24.

16. Agata G, Magdalena F, Karolina O. The application of the Biolog EcoPlate approach in ecotoxicological evaluation of dairy sewage sludge. Appl Biochem Biotechnol. 2014;174(4):1434–43.

17. Paixão SM, Sàágua MC, Tenreiro R,., Anselmo AM. Assessing microbial communities for a metabolic profile similar to activated sludge. Water Environ Res. 2007;79(5):536–46.

18. Zhang TY, Wu YH, Zhuang LL, Wang XX, Hu HY. Screening heterotrophic microalgal strains by using the Biolog method for biofuel production from organic wastewater. Algal Research. 2014;6:175–9.

19. Lopes J, Peixoto V, Coutinho A, Mota C, Fernandes S, editors. Determination of the community-level physiological profiles (CLPP) using BiologTM ECO-plates in the river Lima estuary sediments (Northern Portugal). International Meeting on Marine Research; 2016.

20. Al-Dhabaan FAM, Bakhali AH. Analysis of the bacterial strains using Biolog plates in the contaminated soil from Riyadh community. Saudi J Biol Sci. 2016.

21. Yang L. Ecological restoration and benefit evaluation of the test section of Chaohu Lake wetland (in Chinese). Anhui-Forestry Science and Technology. 2015;41(5):45–7.

22. Cheng Z. The study on ecological restoration modes of wetland landscape based on the natural system: a case study in Chaohu Lake region(in Chinese). Journal of Shenyang Jianzhu University (Social Science). 2015;17(2):122–6.

23. Cao X, Song C, Xiao J, Zhou Y. The Optimal Width and Mechanism of Riparian Buffers for Storm Water Nutrient Removal in the Chinese Eutrophic Lake Chaohu Watershed. Water. 2018;10:1489–99.

24. Li R. Approach to restoration of water environmental ecosystem in Chaohu lake (in Chinese). Journal of Hefei University of Technology (Social Science). 2002;16(5):130–3.

25. Lian Y, Song C, Wu L, Huo L, Cai Z. Study of wetland classification on the north bank of Chaohu based on GIS and RS (in Chinese). Journal of Hefei University of Technology (Social Science). 2008;31(11):1736–9.

26. Wang X, Jiang B, Yang M, Bi X. Ecological and environmental status of Chaohu Lake and protection countermeasures (in Chinese). Yangtze River. 2018;49(17):24–30.

27. Zhang L, Shao S, Liu C, Xu T, Fan C. Forms of Nutrients in Rivers Flowing into Lake Chaohu: A Comparison between Urban and Rural Rivers. Water. 2015;16(3):4523–36.

28. Jones DL, Willett VB. Experimental evaluation of methods to quantify dissolved organic nitrogen (DON) and dissolved organic carbon (DOC) in soil. Soil Biol Biochem. 2006;38(5):991–9.

29. Tate RL. Microbial Communities, Functional versus Structural Approaches. Soil Sci. 1997;163(6):511–2.

30. Sala MM, Arrieta JM, Boras JA, Duarte CM, Vaqué D. The impact of ice melting on bacterioplankton in the Arctic Ocean. Polar Biol. 2010;33(12):1683–94.

31. Garland JL. Analytical approaches to the characterization of samples of microbial communities using patterns of potential C source utilization. Soil Biol Biochem. 1996;28(2):213–21.

32. Yang Y, Yao J, Hua X. Application of rapd in microbial biodiversity identification (in Chinese). J Microbiol. 2000;20(2):23–5.

33. Kong W, Liu K, Liao Z, Zhu Y, Wang B. Effects of organic matters on metabolic functional diversity of soil microbial community under pot incubation conditions (in Chinese). Acta Ecol Sin. 2005;25(9):2291–6.

34. Wang L, Luo X, Peng F, Zhao L. Changes of microbial activity and functional diversity of contaminated soil microbes community in uranium tailings (in Chinese). Environ Sci Technol. 2014;37(3):25–31.

35. Knappová J, Münzbergová Z. Colonization of central European abandoned fields by dry grassland species depends on the species richness of the source habitats: a new approach for measuring habitat isolation. Landscape Ecol. 2012;27(1):97–108.

36. Hemrová L, Münzbergová Z. The effects of plant traits on species’ responses to present and historical patch configurations and patch age. Oikos. 2015;124(4):437–45.

37. Zhou WJ, Sha LQ, Schaefer DA, Zhang YP, Song QH, Tan ZH, et al. Direct effects of litter decomposition on soil dissolved organic carbon and nitrogen in a tropical rainforest. Soil Biol Biochem. 2015;81(6):255–8.

38. Jones DL, Nguyen C, Finlay RD. Carbon flow in the rhizosphere: carbon trading at the soil–root interface. Plant & Soil. 2009;321(1/2):5–33.

39. Hu S, Coleman DC, Carroll CR, Hendrix PF, Beare MH. Labile soil carbon pools in subtropical forest and agricultural ecosystems as influenced by management practices and vegetation types. Agric Ecosyst Environ. 1997;65(1):69–78.

40. Larson WE, Pierce FJ, Dowdy RH. The threat of soil erosion to long-term crop production. Science. 1983;219(4584):458–65.

41. Rasmussen PE, Allmaras RR, Rohde CR, Roager NC. Crop Residue Influences on Soil Carbon and Nitrogen in a Wheat-Fallow System 1. Soil Sci Soc Am J. 1980;44(3):596–600.

42. Rosenstock B, Simon M. Consumption of Dissolved Amino Acids and Carbohydrates by Limnetic Bacterioplankton According to Molecular Weight Fractions and Proportions Bound to Humic Matter. Microb Ecol. 2003;45(4):433–43.

43. Tiquia SM, Davis D,., Hadid H,., Kasparian S,., Ismail M,., Sahly R,., et al. Halophilic and halotolerant bacteria from river waters and shallow groundwater along the Rouge River of southeastern Michigan. Environ Technol. 2007;28(3):297–307.

44. Tiquia SM, Schleibak M,., Schlaff J,., Floyd C,., Benipal B,., Zakhem E,., et al. Microbial community profiling and characterization of some heterotrophic bacterial isolates from river waters and shallow groundwater wells along the Rouge River, southeast Michigan. Environ Technol. 2008;29(6):651–63.

45. Madigan MT, Martinko JM, Parker J. Brock’s Biology of Microorganisms, 9th edn.2000.

46. Shoko I, Chisato T. Effects of dissolved organic matter on toxicity and bioavailability of copper for lettuce sprouts. Environ Int. 2005;31(4):603–8.

47. Lin H, Sun W, Wang F, Wang B, Weng Y, Ma J, et al. Effects of heavy metal within organic fertilizers on the microbial community metabolic profile of a vegetable soil after land application (in Chinese). Journal of Agro-Environment Science. 2016;35(11):2123–30.

48. Li S, Ma D, Zang S, Wang L, Sun H. Structural and functional characteristics of soil microbial community in the Songjiang wetland under different interferences (in Chinese). Acta Ecol Sin. 2018;38(22):7979–89.

49. Yu H. Characteristics of profile soil microbial community structure and function in Poyang Lake wetland (in Chinese). Nanchang: Nanchang University; 2017.

50. Liu J, Wang X, Zhang T, Li X. Assessment of active bacteria metabolizing phenolic acids in the peanut (Arachis hypogaea L.) rhizosphere. Microbiol Res. 2017;205:118–24.

51. Zhou X, Wu F. p-Coumaric acid influenced cucumber rhizosphere soil microbial communities and the growth of Fusarium oxysporum f.sp. cucumerinum Owen. PLoS One. 2012;7(10):e48288.

52. Zhou X, Wu F. Artificially applied vanillic acid changed soil microbial communities in the rhizosphere of cucumber (Cucumis sativus L.). Can J Soil Sci. 2013;93(1):13–21.

53. Preiss J, Romeo T. Molecular Biology and Regulatory Aspects of Glycogen Biosynthesis in Bacteria 1. Prog Nucleic Acid Res Mol Biol. 1994;47:299–329.

54. Subandi S. Role and management of potassium nutrient for food production in Indonesia. Pengembangan Inovasi Pertanian. 2013;6(1):1–10.

55. Meena VS, Maurya BR, Verma JP, Meena RS. Potassium Solubilizing Microorganisms for Sustainable Agriculture2016.

56. Zarjani JK, Aliasgharzad N, Oustan S, Emadi M, Ahmadi A. Isolation and characterization of potassium solubilizing bacteria in some Iranian soils. Archives of Agronomy & Soil Science. 2013;59(12):1713–23.

57. Gundala PB, Chinthala P, Sreenivasulu B. A new facultative alkaliphilic, potassium solubilizing, Bacillus Sp. SVUNM9 isolated from mica cores of Nellore District, Andhra Pradesh, India. Research and Reviews. J Microbiol Biotechnol. 2013;2(1):1–7.

58. Barra Caracciolo A, Bustamante MA, Nogues I, Di Lenola M, Luprano ML, Grenni P. Changes in microbial community structure and functioning of a semiarid soil due to the use of anaerobic digestate derived composts and rosemary plants. Geoderma. 2015;245–246:89–97.

